# Advanced Protection Against Marine Biofouling Using Solar Light

**DOI:** 10.1101/106195

**Authors:** Gabriele Scandura, Rosaria Ciriminna, Lütfiye Yıldız Ozer, Francesco Meneguzzo, Giovanni Palmisano, Mario Pagliaro

## Abstract

We undertook prolonged testing of a new xerogel coating (*AquaSun*) to protect from marine biofouling a surface probe immersed in the seawater of Abu Dhabi. Electron microscopy, Raman mapping, and photoluminescence experiments coupled to extensive analyses of marine water before and after 122 days of testing showed excellent action against biofouling when irradiating the probe with visible light. Considering the high chemical and mechanical stability and the low cost of the sol-gel coating, the technology has significant potential *en route* to replacing conventional antifouling and foul release coatings with a single product of broad applicability.

## Introduction

Every year, more than 80,000 tonnes of marine antifouling (AF) paints, mostly copper-based, are used across the world to protect vessels of any size and scope,^1^ posing a serious threat to the environment. Intense researches carried out at both antifouling coating makers and in academic research centres has actually resulted in the development of several environmentally benign alternatives, mostly based on foul-release polymeric coatings, but including also formulations, whose action is based on less toxic biocides.^2^ Foul-release (FR) waterborne xerogel coatings comprised of organically modified silica (ORMOSIL) are amongst the environmentally benign solutions identified.^3^ Their action is based on the ability to form a thin hydrophobic protective layer to which biofouling sticks loosely to be released even at low cruising speed. Their main limitation is due to their limited antifouling action when the vessel is still in harbor for prolonged periods.

Aiming to develop a coating capable to provide full protection to ships, recreational boats and underwater structures against marine biofouling, we have recently reported the promising discovery that nanoflower-like Bi_2_WO_6_ encapsulated in methyl-modified silica shows excellent photocatalytic antifouling action leading to formation of hydrogen peroxide.^4^ Hydrogen peroxide is a strong and clean oxidant decomposing into water and oxygen,^5^ able to rapidly degrade the (bio)organic species adsorbed onto the film making the surface inhospitable to the settling larvae of fouling organisms, by a free-radical intermediate that prevents the attachment of hard foulants.^6^

Dubbed *AquaSun*, such coating typically forms a transparent thin film (3 µm thick) of methylsilica in which the particles of the semiconductor are homogeneously encapsulated within the ORMOSIL matrix, retaining their flower-like nanostructure. Using uracil as representative molecular precursor of biofouling, we showed that under irradiation with solar light, the coating degraded about half of 1 ppm uracil in three days. No wolframate leaching was observed into the supernatant solution, even after three months immersion, and after repeated irradiation cycles. Remarkably, furthermore, the activity degradation rate was found to be linear, pointing to no saturation effects or prolonged absorption of the substrate. Indeed, further investigation using adenine as probe revealed good foul release properties of the coating, with all adenine adsorbed from a 1 ppm solution being released in 2 hours upon immersion in water, even in the absence of light irradiation. Now, we show the efficiency of this new coating in real life tests conducted for 122 days in sea water withdrawn from Al Raha Beach, Abu Dhabi, United Arab Emirates.

## Results and Discussion

The waterborne coating was sol-gel derived through hydrolytic polycondensation, in acidic conditions, of a sol containing 50 mol % tetraethoxysilane (TEOS) and 50 mol % methyl-triethoxysilane (MTES) to which flower-like nanostructured Bi_2_WO_6_ was added (50 mM) as previously described.^4^ Runs were performed in a 2.5 L beaker, by using sea water (1.5 L) placed in contact with glass slides after getting rid of most sand particles through sedimentation. Bare and functionalized glass slides were used in two different conditions: (**A**) no applied radiation, meaning that only the low diffuse radiation present in the laboratory reached the solution; and (**B**) under intermittent irradiation with 500 W simulated solar light. In the latter case, water evaporation was not negligible. Consequently, an aliquot of sea water (around 100 mL) was poured in the beaker to replenish the original amount around every 10 days since the beginning of the experiment. The overall experiment lasted 122 days.

**Table 1.** Irradiation time under simulated solar light for experiment **B**.

| Period | Hours per day | Number of days | Total hours of irradiation |
| --- | --- | --- | --- |
| From day 1 to day 42 | 2.5 | 42 | 105 |
| From day 43 to day 65 | 8.5 | 23 | 195.5 |
| From day 66 to day 122 | 5 | 57 | 285 |

In experiment **B** the radiation intensities were 2.8 W/m^2^ in the range 315-400 nm and 60 W/m^2^ in the range 450-950 nm. Irradiation was applied every day for the number of hours indicated in Table 1(total number of hours under simulated solar light was 585.5).

Figure 1 and Figure 2 show, respectively, the visible light radiation intensity (wavelength range: 400-1050 nm) in the laboratory during a working day, and the visible light irradiation intensity (wavelength range: 400-1050 nm) in the laboratory during a public holiday. Radiation intensity was measured with a Delta Ohm 9721 radiometer and the matching probes.

**Figure 1.**
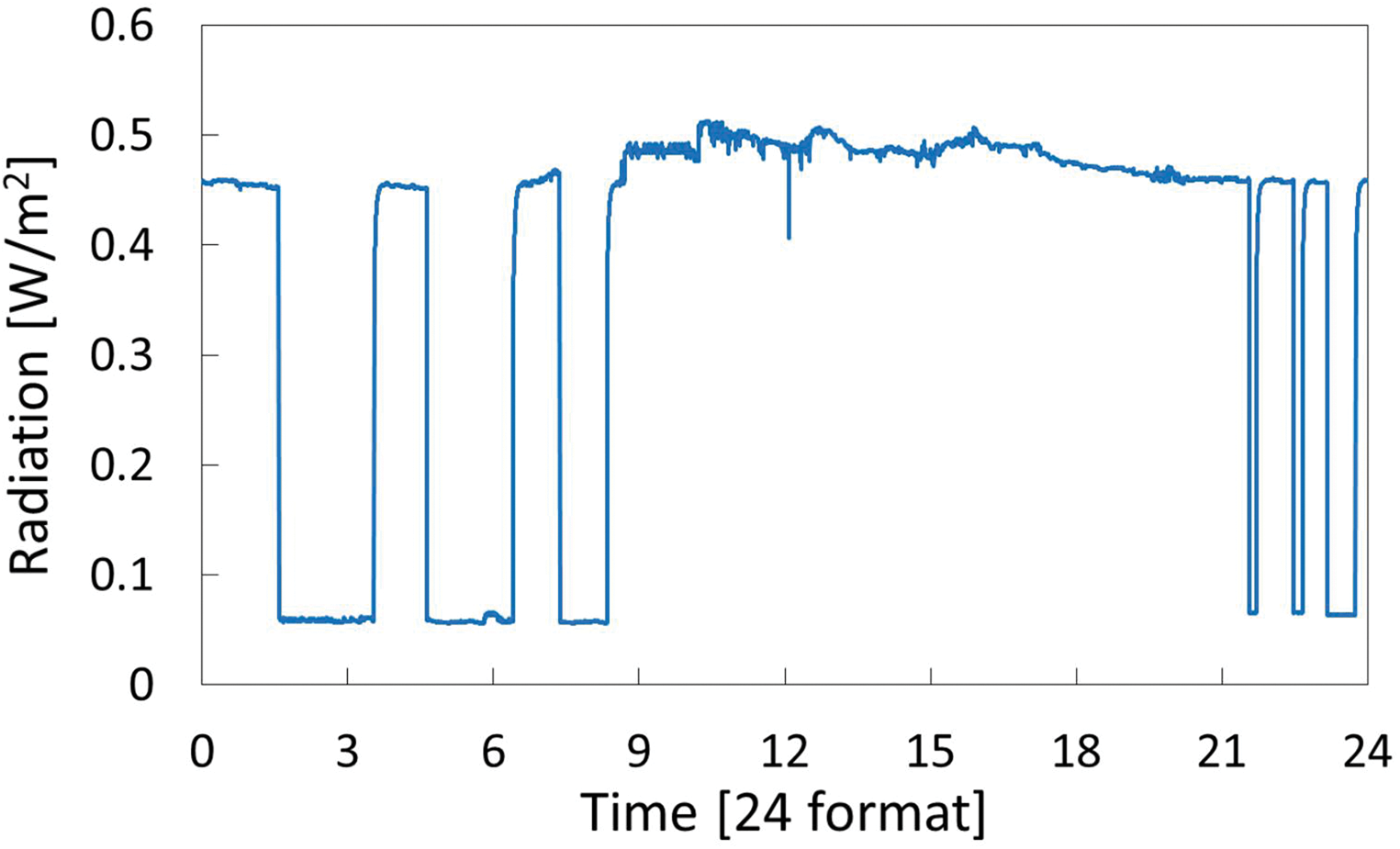
Radiation intensity (wavelength range: 400-1050 nm) in the laboratory during a working day.

**Figure 2.**
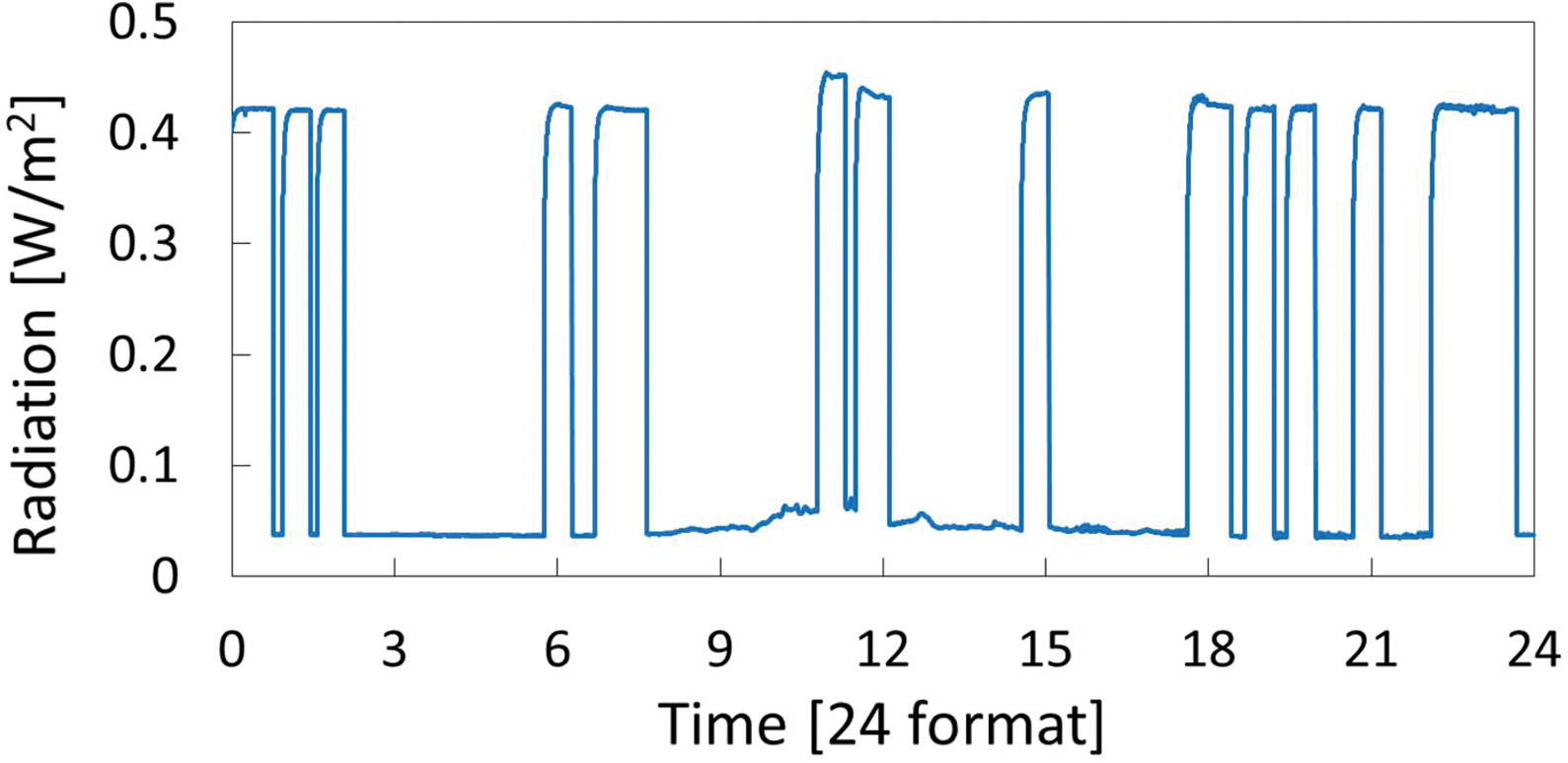
Radiation intensity (wavelength range: 400-1050 nm) in the laboratory during a public holiday.

Characteristics of the sea water were analyzed with a Shimadzu TOC-L Total Organic Carbon analyzer, a Delta Ohm pH-χ-O_2_ meter (mod. HD22569.2), an Ion Chromatography System Dionex ICS-5000 and a HACH 2100AN turbidimeter. Table 2 and Table 3 summarize the characteristics of the sea water at the beginning (day 0) and at the end (day 122) of the experiments.

**Table 2.** Main characteristics of the sea water at the beginning (day = 0) and at the end (day = 122) of the experiment.

|  | Day 0 | Day 122 experiment A | Day 122 experiment B |
| --- | --- | --- | --- |
| pH | 8.250 | 8.483 | 8.191 |
| Conductivity [mS] | 50.4 | 53.6 | 64.5 |
| Dissolved O<sub>2</sub> [mg/L] | 7.73 | 7.53 | 7.43 |
| TC [mg/L] | 26.60 | 26.76 | 25.72 |
| IC [mg/L] | 25.38 | 25.57 | 22.93 |
| TOC [mg/L] | 1.218 | 1.186 | 2.784 |
| Turbidity [NTU] | 0.130 | 0.226 | 0.407 |

TC=total carbon, IC=inorganic carbon, TOC=total organic carbon

**Table 3.**
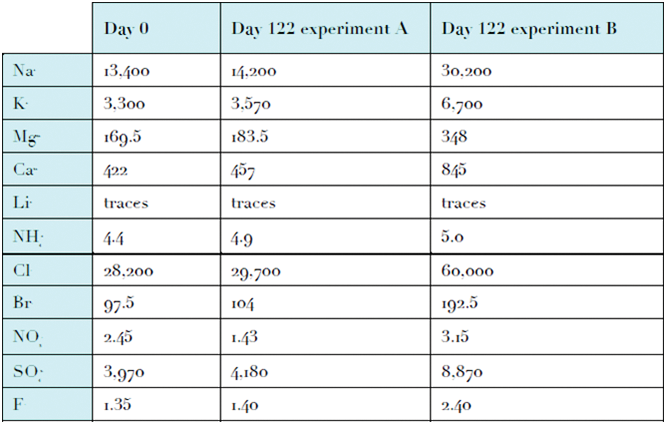
Ions concentrations expressed in mg/L in the sea water at the beginning (day = 0) and at the end (day = 122) of the experiment.

|  | Day 0 | Day 122 experiment A | Day 122 experiment B |
| --- | --- | --- | --- |
| Na | 13,400 | 14,200 | 30,200 |
| K | 3,300 | 3,570 | 6,700 |
| Mg | 169.5 | 183.5 | 348 |
| Ca | 422 | 457 | 845 |
| Li | traces | traces | traces |
| NH<sub>4</sub><sup>+</sup> | 4.4 | 4.9 | 5.0 |
| Cl | 28,200 | 29,700 | 60,000 |
| Br | 97.5 | 104 | 192.5 |
| NO<sub>3</sub><sup>-</sup> | 2.45 | 1.43 | 3.15 |
| SO<sub>4</sub><sup>2-</sup> | 3,970 | 4,180 | 8,870 |
| F | 1.35 | 1.40 | 2.40 |

Under solar light, marine phytoplankton microorganisms give place to photosynthesis providing organic matter for the organisms that comprise the majority of marine life,^7^ consuming inorganic carbon. For this reason, in the experiment **B** the total organic carbon (TOC) increased and the inorganic carbon (IC) decreased when comparing the initial and final concentrations. Nevertheless, the total carbon (TC) is reduced because the thin film under visible light is able to mineralize organic matter with formation of carbon dioxide which evolves from the supernatant solution.^1^

Conversely, in experiment **A**, the values of TC, IC, and TOC before and after each run are substantially the same, which is due to the insufficient radiation, thereby proving that degradation is due to the photocatalytically formed hydrogen peroxide achievable only by irradiating with (simulated) solar light. When the film is applied on a surface constantly exposed to solar radiation, it continuously produces H_2_O_2_ according to Equations 1–5, where h^+^ are the holes generated in the valence band of the semiconductor and e^-^ are the electrons formed in the conduction band:^8^

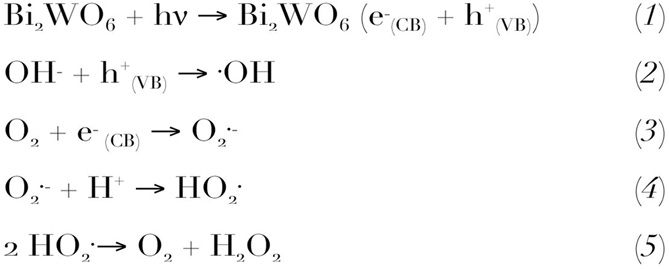

Diffuse reflectance spectra were measured by UV-Vis spectrophotometer (Shimadzu UV-2600) in the range 350-750 nm. Comparison between the UV-vis diffuse reflectance spectra of the bare glass slide and the glass slide functionalized with the *AquaSun* coating (Figure 3) highlights a minor shift of the latter since, under simulated solar light, the film functionalized with the coating is much cleaner than the unprotected glass.

**Figure 3.**
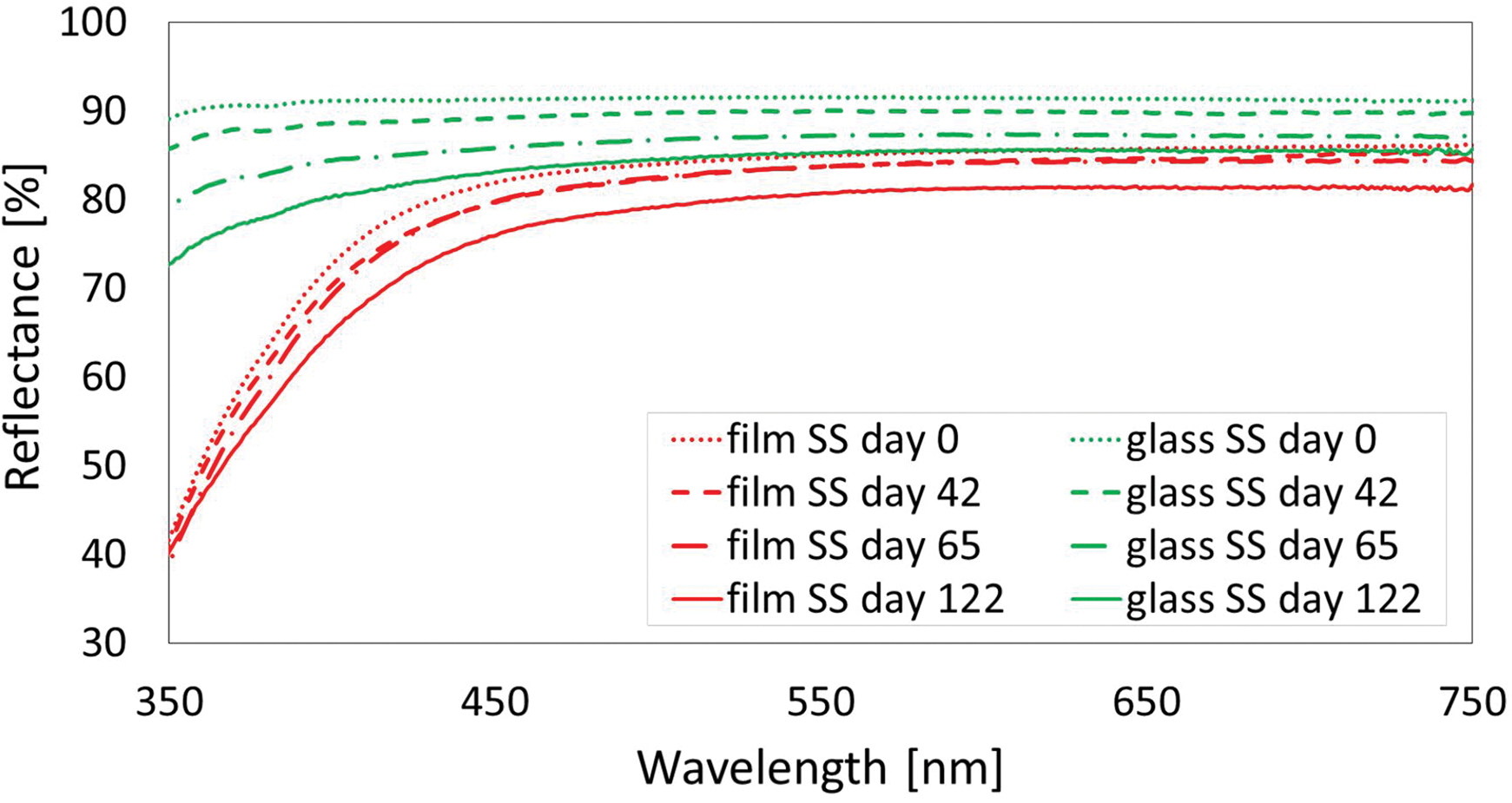
Diffuse reflectance spectra of bare glass and functionalized glass (labelled as “film”) in the presence of simulated solar light. SS stands for “solar simulator” and it refers to the experiment **B**.

**Figure 4.**
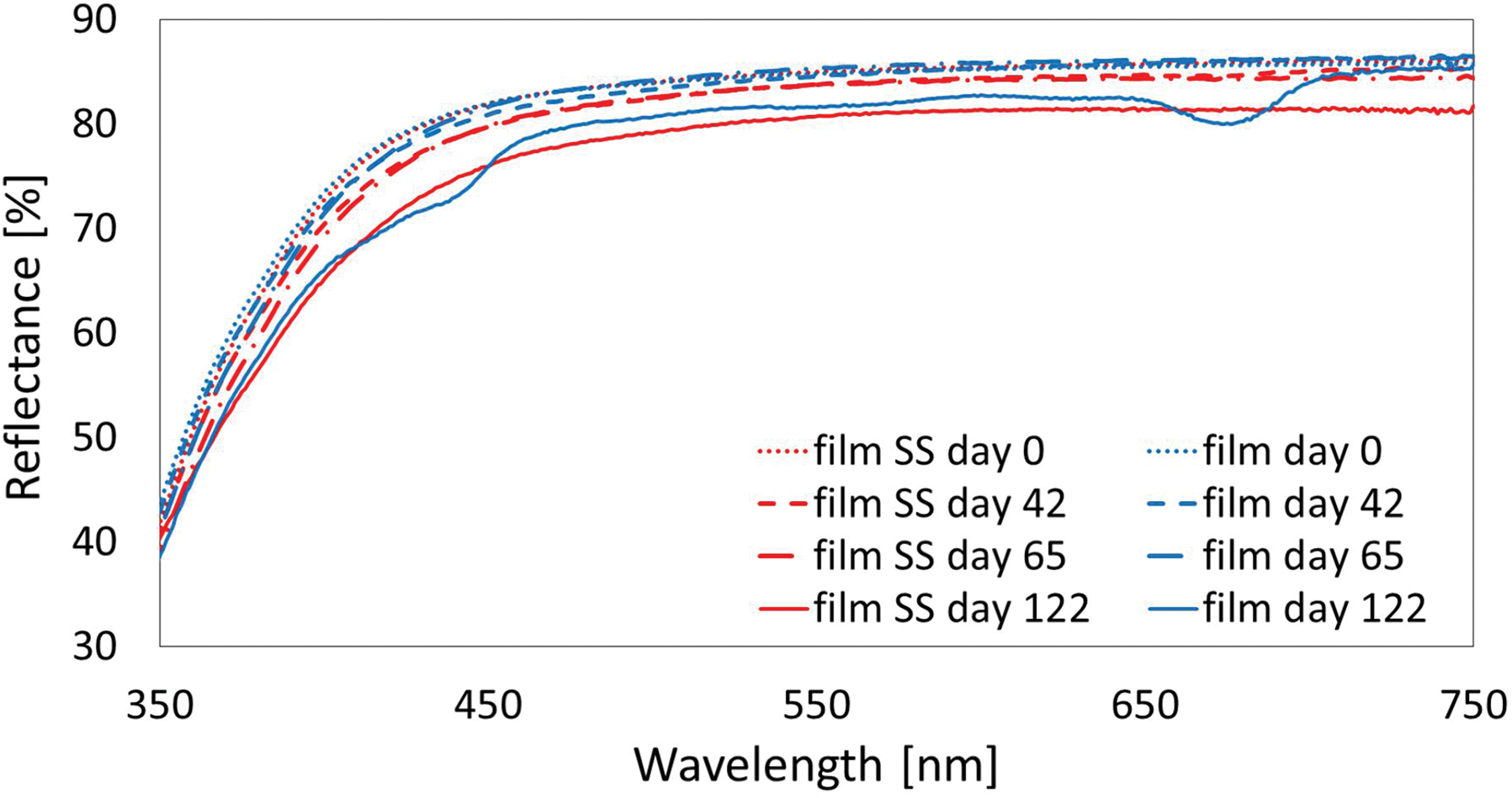
Diffuse reflectance spectra of functionalized glass (labelled as “film”) in the absence or in the presence of simulated solar light. SS stands for “solar simulator” and it indicates the experiment **B**.

The effect of the solar light on the glass slide functionalized with the *AquaSun* thin film is evident from Figure 4 showing that the coated glass, when exposed to the low diffuse radiation present in the laboratory room, undergoes the growth of biomass on its surface. because after 4 months (day 122) two significant peaks (around 440 and 675 nm) appear in the diffuse reflectance spectrum. Accordingly, the TC, IC and TOC values at the end of the test in sea water did not decrease: a crucially important result that was visually confirmed (see photograph in Figure 5).

**Figure 5.**
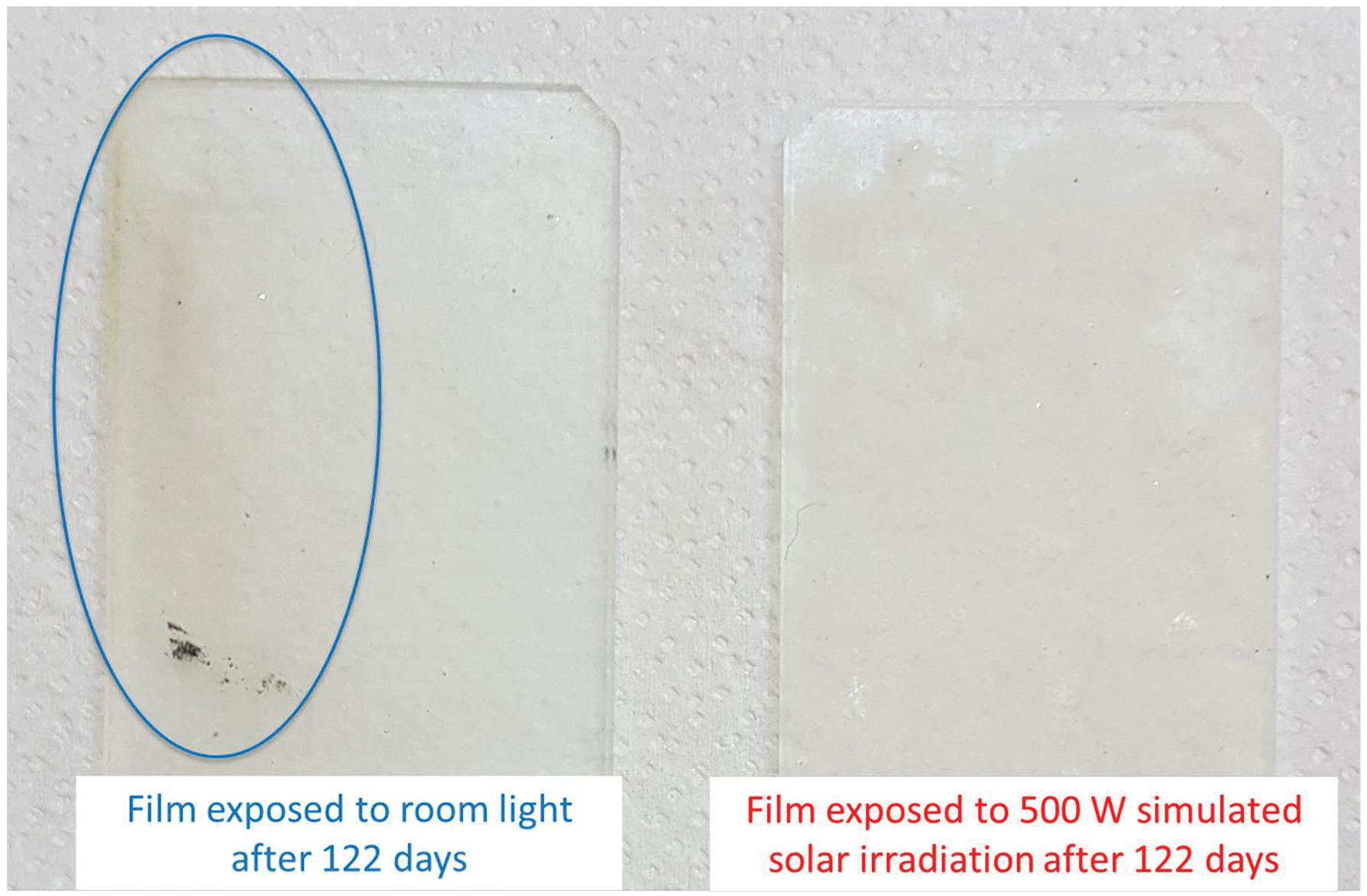
The blue oval indicates the area of the film where a biomass layer deposited because of the low irradiation.

The presence of the biomass also affects the photoluminescence (PL) of the film. The spectra in emission mode with an excitation at 300 nm were recorded using a Perkin Elmer LS 55 spectrometer between 310 and 600 nm (200 nm/min scan rate). The PL spectrum of the functionalized glass shows a peak around 420 nm (Figure 6) which is clearly stronger when the film was exposed to room light in comparison to the peak shown by the coated glass exposed to simulated solar irradiation. Indeed, it is known that some marine microorganisms such as algae can produce photoluminescence emission in the visible region due to the proteins or aromatic amino acid, and their metabolites.^9^

**Figure 6.**
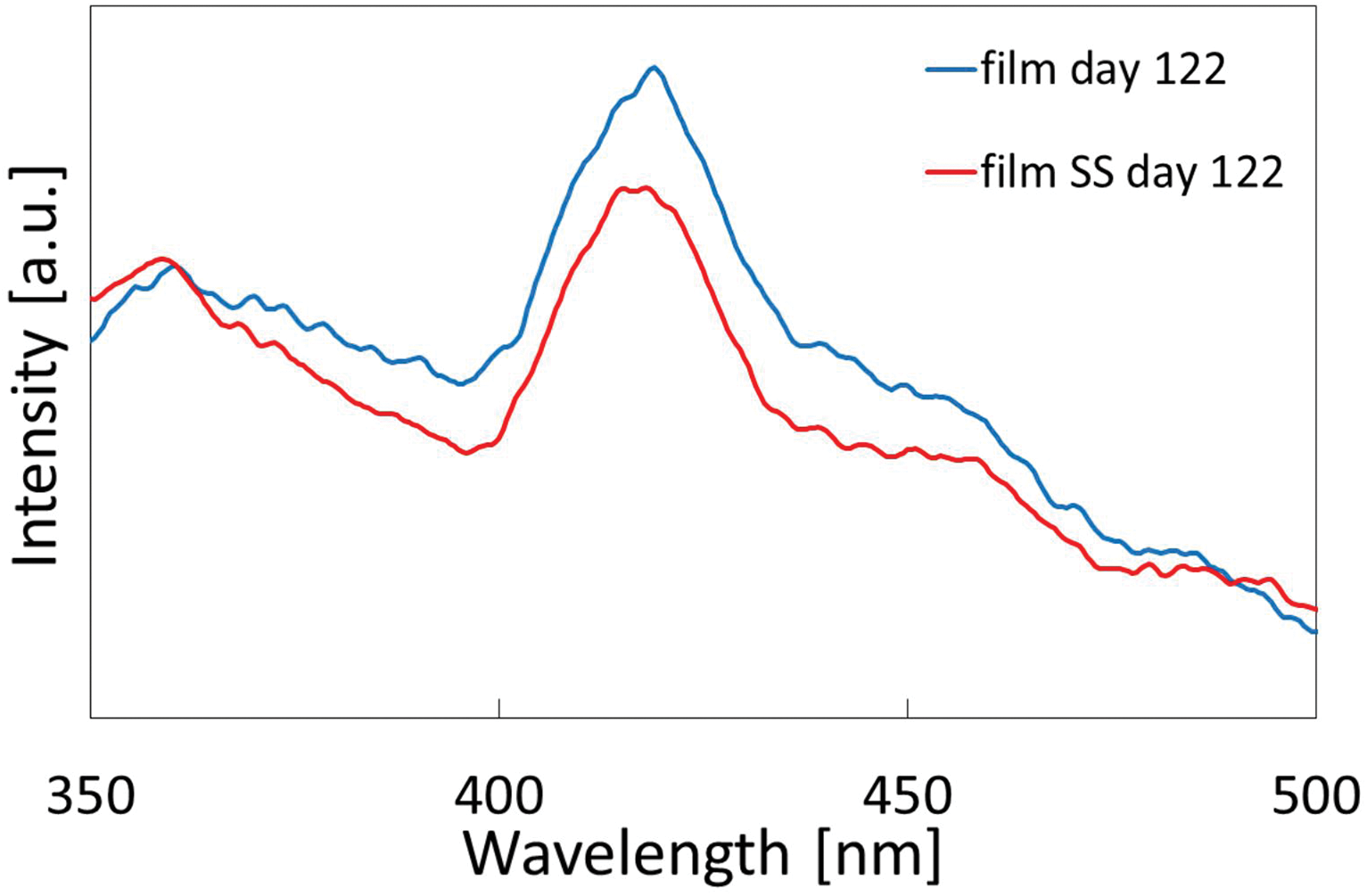
The photoluminescence (PL) spectra of functionalized glass (labelled as “film”) with an excitation of 300 nm in the absence or in the presence of simulated solar light. “SS” stands for “solar simulator” and it refers to experiments **B**.

Figure 7 depicts the Raman spectra of both bare and functionalized glass before (day 0) and after the experiments (day 122), recorded with a Witec Alpha 300R equipment, with an excitation wavelength of 532 nm and a laser power of ca. 75 mW. Scans were taken over an extended range (100-2000 cm−^1^) with 5 s integration time and 30 accumulations. Raman mapping was done with 0.1 s integration time.

**Figure 7.**
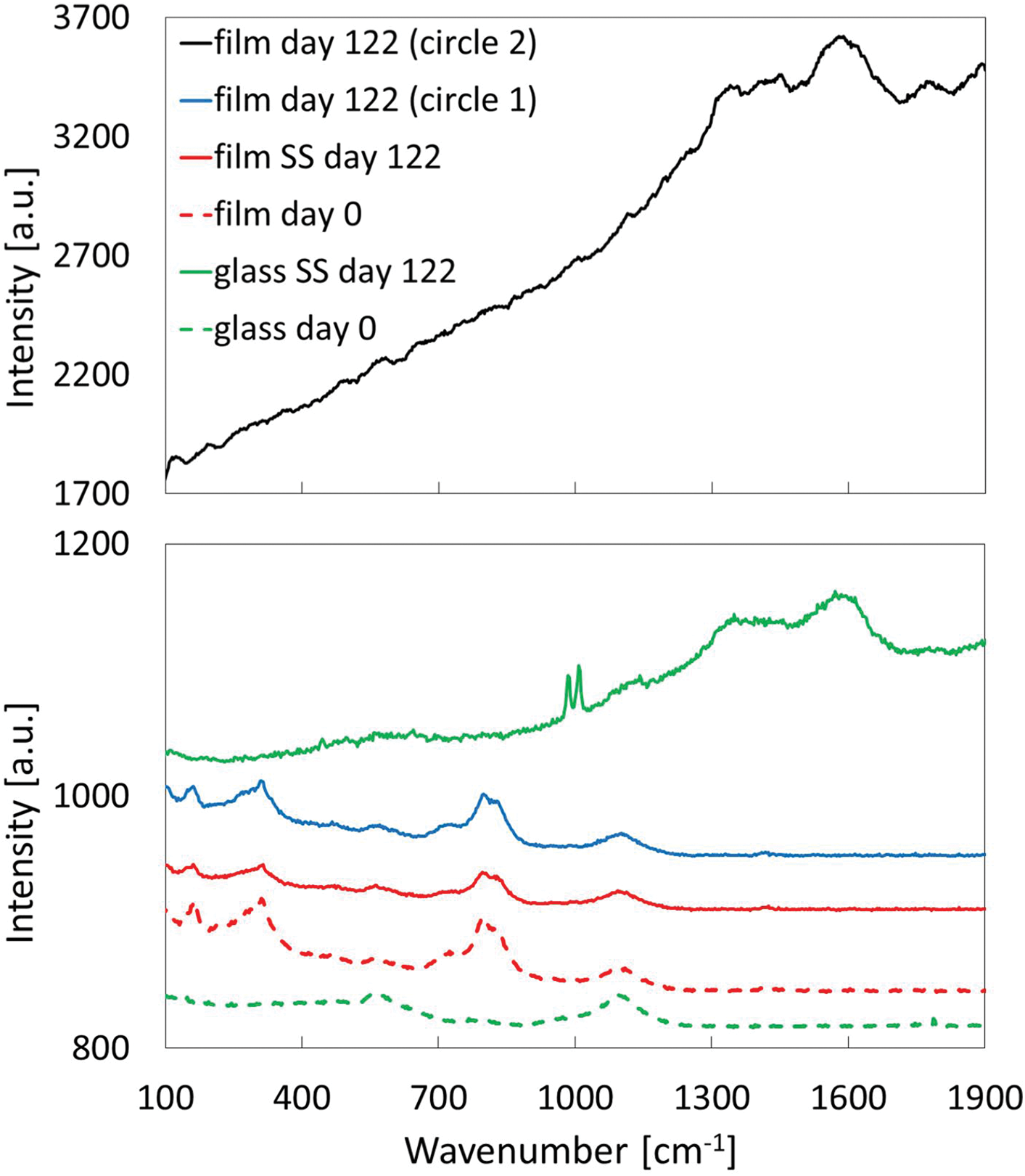
Raman spectra of functionalized (labelled as “film”) and bare glass before (day 0) and after 122 days in the absence or in the presence of simulated solar light. “SS” stands for “solar simulator” and it refers to experiments B. The curves related to the ‘film day 122’ were obtained by pointing Raman laser on the circles 1 and 2 in Figure 8a.

The glass coated with *AquaSun* coating exhibits three peaks (at ca. 161, 311, and cm−^1^) attributable to Bi_2_WO_6_ encapsulated in the ORMOSIL,^10^ and two peaks (around 560 and 1095 cm−^1^) due to the glass substrate.

In experiment **B**, the functionalized film after four months shows the same Raman spectrum as day = 0. On the contrary the bare glass spectrum displays several new peaks. The film of the experiment **A** affords two different types of spectra according to where the laser beam is pointed. One (corresponding to circle 1 in Figure 8a) is identical to the spectrum at day = 0. Another (corresponding to circle 2 in Figure 8a) displays a much larger background intensity (top curve in Figure 7). This spectrum results from biomass which has been produced because of the poor illumination in the room, and is caused by the photoluminescence of the biomass itself (in agreement with Figure 6), which covers completely the signals of Bi_2_WO_6_.

**Figure 8.**
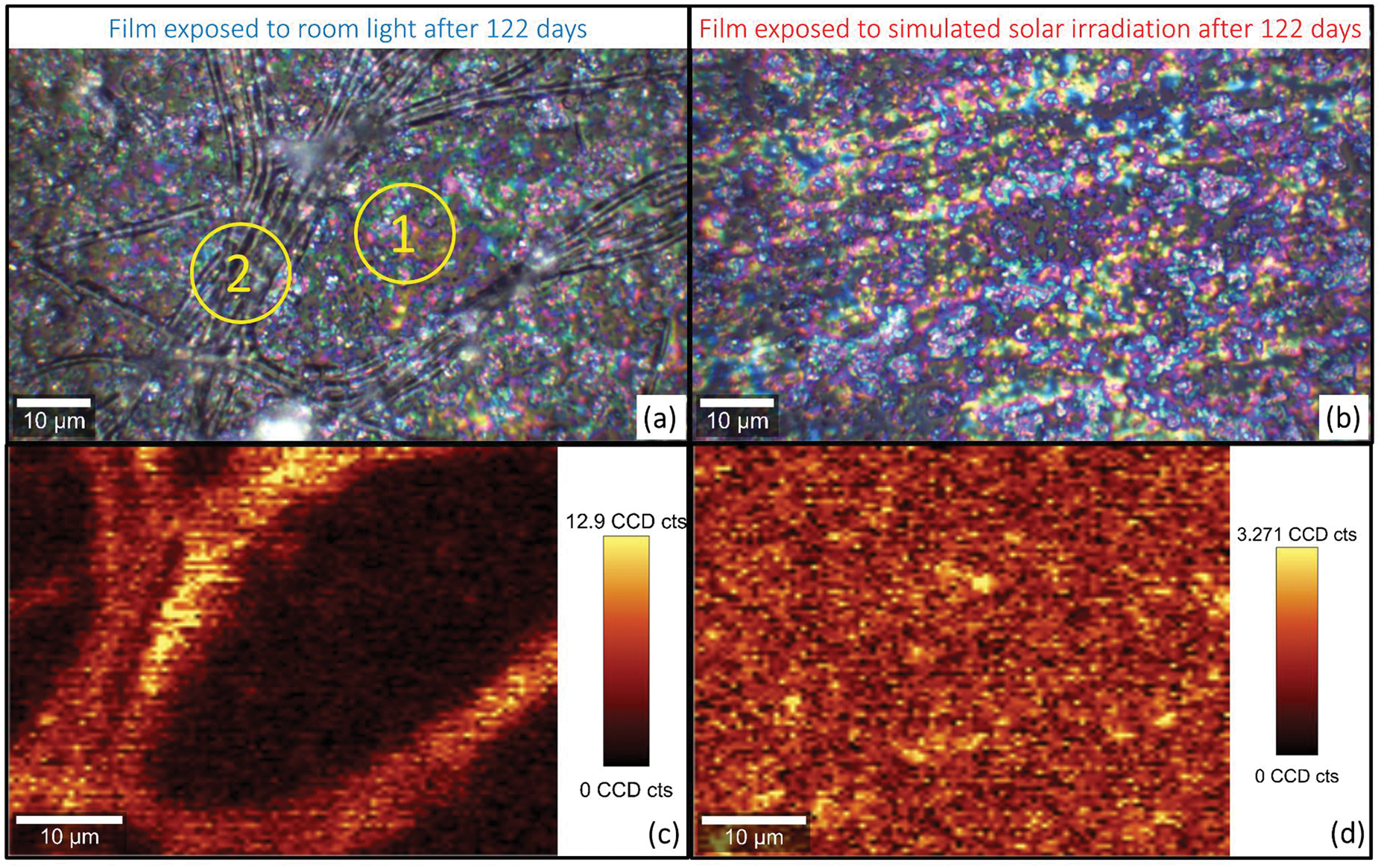
100x magnified optical images of the films after 122 days in the absence (a) or in the presence of simulated solar light (b). 799 cm-1 peak Raman mapping (c, d) corresponding to the images above respectively. Circles 1 and 2 indicate where the laser has been pointed during the recording of the curves labelled as ‘film day 122’ in Figure 7.

Remarkably, these two populations of spectra obtained from the film irradiated by room light only are clearly distinguishable in Figure 8c, where the Raman mapping of 799 cm ^-1^ peak is displayed: the area where the biomass is abundant gives a stronger intensity, and this is a further confirmation that the far more intense signal of the black curve in Figure 7 is due to the biomass. On the other hand, the Raman mapping of the film exposed to solar light indicates a homogeneous surface, highlighting the absence of biomass in that surface, with a Raman spectrum which does not change with the area analysed, and a signal intensity far lower than the one given by the film with accumulated biomass.

The conclusive evidence that *AquaSun* inhibits the formation of biomass on the surface faced to solar light is given by SEM analysis (Figure 9, carried out with a FEI Nova NanoSEM 650 microscope). No relevant differences are observed in the glass coated with *AquaSun* at day = 0 and after 122 days under the conditions of experiment **B** (Figures 9a and 9b, respectively). Conversely, the film exposed to room light after 122 days is significantly populated by biomass (Figure 9c). As with regards to the bare (unprotected) glass in experiment **B**, the SEM image (figure 9d and inset) demonstrates the presence of microorganisms with a circular shape, closely resembling a planktonic diatom.

**Figure 9.**
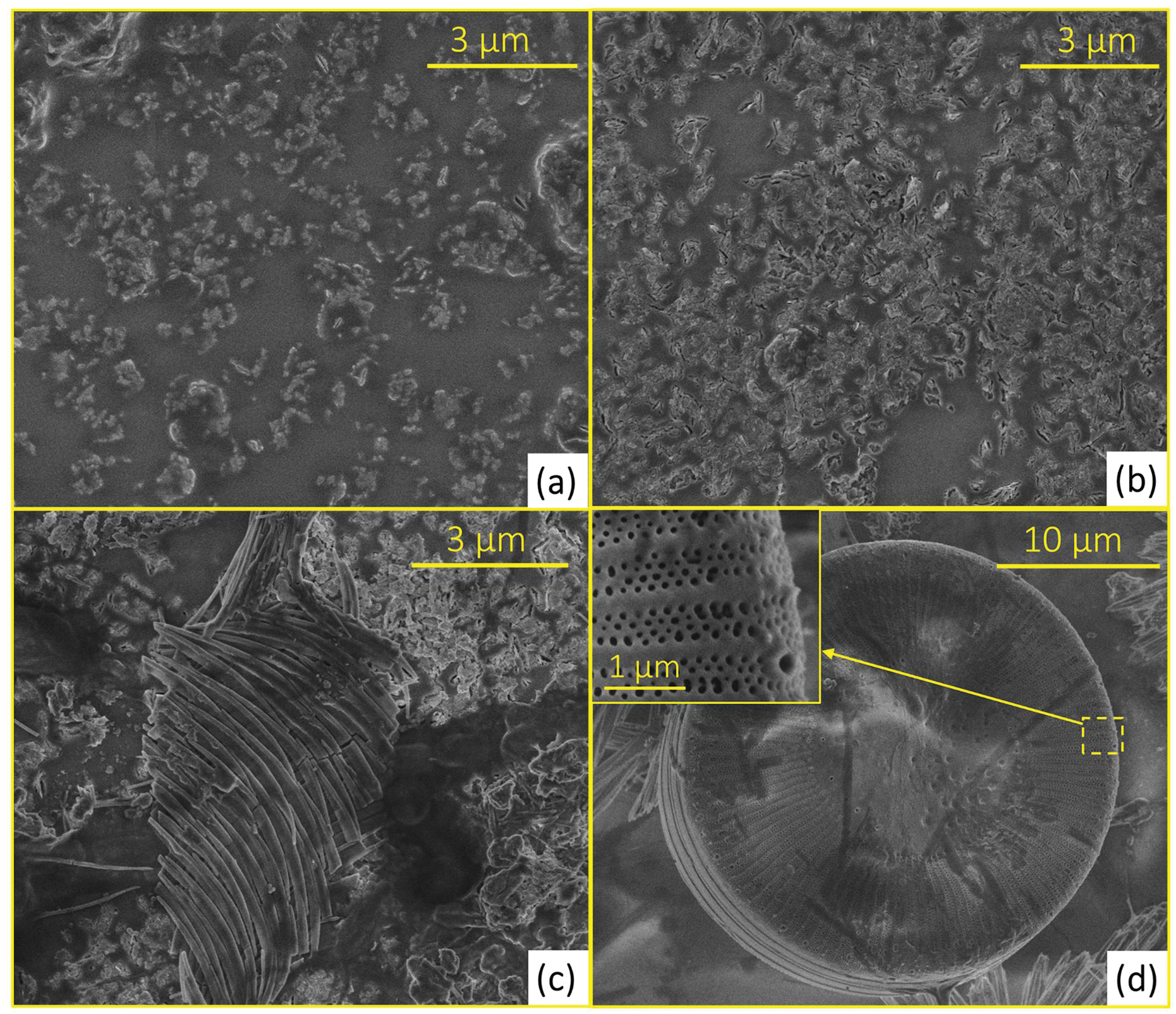
SEM images at 25000x magnification of the functionalized glass at day 0 (a), after experiment **B** (b) and after experiment **A** (c); SEM image at 8000x magnification of the bare glass after experiment **B** (d).

It is worth reminding that the photocatalytic reaction on the submarine surface is triggered by visible-light radiation which easily penetrates sea water, with reflection at moderate depths accounting to only 5-10 %, and with a negligible absorption.^11^

To assist in the process for identifying alternatives to be used as substitutes to replace copper-based antifouling technology, which is of high environmental concern, we provide a brief insight on how the use of *AquaSun* ensures that copper is not being replaced with equal or more hazardous alternative, thereby getting ahead of new bans or restrictions. For example, phase out of copper-based anti-fouling paints on recreational vessels less than 65 feet in length by 2020 was adopted by the State of Washington, where salmon fisheries generate $3.18 billion of economic activity each year supporting over 17,000 jobs, with even 2 ppb levels of copper interfering with a salmon’s its ability to avoid predators and also adversely impacts the ability of fish eggs and fry to develop normally.^12^

The components of the new antifouling product are Bi_2_WO_6_ and the encapsulant methylisica matrix. The only acute effect recorded in Material Safety Data Sheet for Bi_2_WO_6_ is moderate irritation of the respiratory system following inhalation, and low toxicity following ingestion due to its insoluble nature. The complete lack of solubility in water is most promising for what concerns its aquatic toxicity and ecotoxicity (persistance in the environment and bioaccumulation).

In general, bismuth is a heavy metal element with unusual low toxicity, obtained as a byproduct of extraction of other metals, including tungsten, used in many common stomach remedies and cosmetic products.^13^ Tungsten, in its turn, does not constitute an important health hazard.^14^ The tungstate anion WO_4_^2^− isomorph to molybdate, antagonizes the normal metabolic action of isomorph molybdate MoO_4_^2^− as metal carrier. Finally, the low cost, facile preparation with high purity, high stability in various electrolytes, and resistance to photocorrosion make Bi_2_WO_6_ ideally suited for application to photocatalytic wastewater treatment.^15^

Sol-gel derived methylsilica is an eminent ORMOSIL. Due to their inertness and excellent textural properties, these materials are used as carriers for drugs and bioactive agents,^16^ with ORMOSIL nanoparticles generally being of little or no toxicity.^17^ Methyl-modified silica is highly stable in water and even more in seawater, whose pH is limited to the range 7.5 to 8.4. Similar methyl-modified silica coatings functionalized with silver ions are being considered for as antibacterial coatings on surgical-grade stainless steel.^18^

Being a waterborne xerogel organosilica coating,^19^ *AquaSun* is easily applied to all type of surfaces (steel, fiberglass, aluminum, wood etc.) by spraying or brushing eliminating the need to handle dangerous paint formulations (safe for workers), curing at room temperature. The high chemical and physical stability of antifouling sol-gel coatings generally affords minimum of two years protection, thereby reducing frequency of applications, while reducing drag to the smooth layer formed at the outer surface of the vessel. Forthcoming trials in the open sea will be aimed to identify the optimal application parameters for different vessels. The technology, indeed, is made available in open innovation regime with the aim to shorten time to the introduction of high performance, environmentally friendly antifouling coatings affording affording the economic advantages of efficient antifouling action, while protecting the aquatic environment and, ultimately, human health.

## Conclusions

The use of Bi_2_WO_6_ encapsulated in ORMOSIL (*Aquasun*) and deposited on glass has been proven effective in preventing the accumulation of biomass on glass in a real sea water environment in prolonged testing lasting up to four months. Extensive Raman mapping characterization of the coating using a glass surface as probe highlights the formation of biomass on selected areas of the glass when only diffuse visible light radiation reaches the surface, whereas unprotected glass was found covered with diatoms. Exposing the coating to visible light radiation using simulated solar light even for a few hours per day (2.5, 5 and 8 h for 122 consecutive days) at moderate UV-vis radiation (2.8 and 60 W/m^2^ in the UV and visible regions, respectively), prevents the formation of any biofilm with a significant reduction in the total amounts of inorganic carbon in water and a concomitant increase of dissolved organic carbon due to ongoing photosynthesis. These results open the route to widespread application of the *AquaSun* for the protection of widely different surfaces constantly submerged in sea water.

## Acknowledgments

This article is dedicated to Dr. Valerica Pandarus (SiliCiycle, Canada) on the joint occasion of 10 years of fruitful cooperation with two of us (RC and MP) and of her PhD from Laval University. Shadi W. Hasan (Masdar Institute of Science and Technology, Abu Dhabi) is acknowledged for enabling the use of the turbidimeter in his lab. We are grateful to Cyril Aubry (instructor at Masdar Institute of Science and Technology) for his precious assistance during SEM and Raman measurements.

## References

1. D. Williams, Challenges in developing antifouling coatings, IMarEST, London, 29 April 2010.

2. R. Ciriminna, F. V. Bright, M. Pagliaro, Ecofriendly Antifouling Marine Coatings, ACS Sustainable Chem. Eng. 2015, 3, 559–565.

3. M. R. Detty, R. Ciriminna, F. V. Bright, M. Pagliaro, Environmentally Benign Sol-Gel Antifouling and Foul-Releasing Coatings, Acc. Chem. Res. 2014, 47, 678–687.

4. G. Scandura, R. Ciriminna, Y.-J. Xu, M. Pagliaro, G. Palmisano, Nanoflower-Like Bi2WO6 Encapsulated in ORMOSIL as a Novel Photocatalytic Antifouling and Foul-Release Coating, Chem. Eur. J. 2016, 22, 7063–7067.

5. R. Ciriminna, L. Albanese, F. Meneguzzo, M. Pagliaro, Hydrogen Peroxide: A Key Chemical for Today's Sustainable Development, ChemSusChem 2016, 9, 3374–3381.

6. S. Møller Olsen, J. B. Kristensen, B. S. Laursen, L. T. Pedersen, K. Dam-Johansen, S. Kiil, Antifouling effect of hydrogen peroxide release from enzymatic marine coatings: Exposure testing under equatorial and Mediterranean conditions, Prog. Org. Coat. 2010, 68, 248–257.

7. P. Falkowski, Ocean science: The power of plankton, Nature 2012, 483, S17–S20.

8. N. Zhang, R. Ciriminna, M. Pagliaro, Y.-J. Xu, Nanochemistry-derived Bi_2_WO_6_ nanostructures: towards production of sustainable chemicals and fuels induced by visible light, Chem. Soc. Rev. 2014, 43, 5276–5287.

9. S. Determann, J. M. Lobbes, R. Reuter, J. Rullkötter, Ultraviolet fluorescence excitation and emission spectroscopy of marine algae and bacteria, Mar. Chem. 1998, 62, 137–156.

10. A. Phuruangrat, P. Dumrongrojthanath, N. Ekthammathat, S. Thongtem, T. Thongtem, Hydrothermal Synthesis, Characterization, and Visible Light-Driven Photocatalytic Properties of Bi2WO6 Nanoplates, J. Nanomater. 2014, 1 (2014) Article ID 138561.

11. D. M. McMaster, S. M. Bennett, Y. Tang, J. A. Finlay, G. L. Kowalke, B. Nedved, F. V. Bright, M. E. Callow, J. A. Callow, D. E. Wendt, M. G. Hadfield, M. R. Detty, Antifouling character of ‘active’ hybrid xerogel coatings with sequestered catalysts for the activation of hydrogen peroxide, Biofouling 2009, 25, 21–33.

12. Washington State Legislature, Chapter 70.300 Recreational Water Vessels, Antifouling Paints, http://apps.leg.wa.gov/Rcw/default.aspx?cite=70.300&full=true

13. J. Krüger, P. Winkler, E. Lüderitz, M. Lück, H. U. Wolf, Bismuth, Bismuth Alloys, and Bismuth Compounds, Ullmann’s Encyclopedia of Industrial Chemistry, Wiley-VCH, Weinheim: 2003.

14. P. E. Leffler, G. Kazantzis, “Tungsten” In Handbook on the Toxicology of Metals, G. F. Nordberg, B. A. Fowler, M. Nordberg (Ed.s), 4^th^ edition, Elsevier: 2016; Chapter 58.

15. S. Girish Kumar, K. S. R. Koteswara Rao, Tungsten-based nanomaterials (WO_3_ & Bi_2_WO_6_): Modifications related to charge carrier transfer mechanisms and photocatalytic applications Appl. Surf. Sci. 2015, 355, 939–958.

16. I. Roy, P. Kumar, R. Kumar, T. Y. Ohulchanskyy, K.-T. Yong, P. N. Prasad, Ormosil nanoparticles as a sustained-release drug delivery vehicle, RSC Adv. 2014, 4, 53498–53504.

17. D. Kumar, C. K. Prashant, A. Kumar Dind, S. Mitra, “Toxicity assessment of Ormosil Nanoparticles” In Nano BioMaterials, V. Rajendran, P. Prabu, K. E. Geckeler (Ed.s), Bloomsbury Publishing India, New Delhi: 2014.

18. R. Procaccini, A. Bouchet, J. I. Pastore, C. Studdert, S. Ceré, S. Pellice, Silver-functionalized methyl-silica hybrid materials as antibacterial coatings on surgical-grade stainless steel, Prog. Org. Coat. 2016, 97, 28–36.

19. M. R. Detty, R. Ciriminna, F. V. Bright, M. Pagliaro, Xerogel Coatings Produced by the Sol–Gel Process as Anti-Fouling, Fouling-Release Surfaces: From Lab Bench to Commercial Reality, ChemNanoMat 2015, 1, 148–154.

